# Insulin-like androgenic gland factor regulates the development of internal and external secondary sexual characteristics in juvenile male kuruma prawn *Marsupenaeus japonicus*

**DOI:** 10.1101/2025.05.29.656773

**Authors:** Takehiro Furukawa, Fumihiro Yamane, Rino Takeuchi, Tsuyoshi Ohira, Kenji Toyota, Taeko Miyazaki, Naoaki Tsutsui

## Abstract

In decapod crustaceans, insulin-like androgenic gland factor (IAG) is a well-known peptide hormone produced by the androgenic gland, a male-specific endocrine organ. In certain species classified within Pleocyemata, experimental manipulation of IAG signaling has successfully altered both internal and external sexual characteristics, leading to sex reversal. Therefore, IAG is considered a key regulator of male sexual differentiation and maturation and is thought to be a crustacean androgen. In the kuruma prawn *Marsupenaeus japonicus* (Dendrobranchiata), an important fishery species in Japan, a homologous molecule known as Maj-IAG has been identified. However, its physiological function in males remains poorly understood. In this context, we conducted *Maj-IAG* knockdown (KD) via RNA interference in juvenile male *M*. *japonicus*. Over a 9-week period, *Maj-IAG* KD significantly inhibited the development of the petasma, male-specific external copulatory structure, and parts of the internal reproductive organs, such as the vas deferens and seminal vesicles. In contrast, somatic growth and testicular development were unaffected by *Maj-IAG* KD. Histological analysis of the testes revealed no evident abnormalities such as degeneration of testicular structures or abnormal arrest of germ cell differentiation. These findings indicate that Maj-IAG partially governs the development of male secondary sexual characteristics in juveniles and further suggest that once testicular differentiation is initiated, the maintenance and development of the testis proceeds without dependence on *Maj-IAG*. Our study provides fundamental insights that may contribute to the advancement of functional sex reversal technologies in penaeid shrimps.

**Highlights:**

- Long-term gene knockdown of the insulin-like androgenic gland factor gene (*Maj-IAG*) was successfully achieved in *Marsupenaeus japonicus*.
- Somatic growth and spermatogenesis were not affected by *Maj-IAG* knockdown.
- *Maj-IAG* is involved in the development of male secondary sexual characteristics in juvenile prawns, including the external male copulatory structures and certain parts of the internal male reproductive organs.

## 1. Introduction

The global demand for commercially valuable fishery species is increasing. Owing to sex-related differences in growth characteristics and economic value, research on sex-control technologies in target species for aquaculture and their potential commercial applications is advancing. For example, in Nile tilapia *Oreochromis niloticus*, males grow larger and faster than females (Mair et al., 1995), and large-scale production of all-male populations has been achieved by inducing genetic females to develop functional male phenotypes through sex steroid hormone treatment, thereby improving productivity (Mair et al., 1997). In the case of hybrid sturgeon (*Huso huso* × *Acipenser ruthenus*), studies have been conducted on sex reversal using sex steroid hormones and related technologies to enhance the efficiency of caviar production (Kinami and Ineno, 2025; Omoto et al., 2002). In the tiger pufferfish *Takifugu rubripes*, the testes have high economic value as food. Therefore, a combination of hormonal sex reversal and genetic sexing methods has been established (Matsunaga et al., 2014; Rashid et al., 2007), enabling the efficient and reliable production of all-male populations. Hormonal sex reversal techniques, which are often used in conjunction with genetic sex identification, have been applied in commercial aquaculture.

In decapod crustaceans, sexual dimorphism in growth is well-known, and the development of monosex populations with superior growth performance is considered advantageous for aquaculture. Insulin-like androgenic gland factor (IAG) has received the most attention as a primary factor in the production of monosex population (Levy and Sagi, 2020). IAG is a peptide hormone predominantly produced by the androgenic gland (AG), a male-specific endocrine organ unique to malacostracan crustaceans. Manipulation of IAG signaling has been shown to induce sex reversal in some species. For example, in the giant freshwater prawn *Macrobrachium rosenbergii*, successful bidirectional sex reversal, both male-to-female and female-to-male, has been achieved through *IAG* silencing, surgical AG ablation, and surgical AG implantation (Malecha et al., 1992; Nagamine et al., 1980; Ventura et al., 2012). In male-to-female sex reversal, suppression of the development of male-specific external structures, such as the gonopores and appendices masculinae, along with the induction of oogenesis, has been observed. In the white pacific shrimp *Litopenaeus vannamei*, functional sex reversal has not been confirmed to date; however, AG ablation results in impaired petasma development as well as reduced somatic growth (Alfaro-Montoya et al., 2016). Similarly, in the red swamp crayfish *Procambarus clarkii*, although no feminization was observed, the development of the first abdominal appendages, a secondary sexual characteristic, and spermatogenesis in the testes were suppressed following *IAG* silencing (Ge et al., 2020). Collectively, these findings support the role of IAG as a key regulator of male sexual differentiation and maturation in decapod crustaceans. Levy et al. (2018) proposed the concept of the “IAG-switch” as a universal regulatory mechanism underlying sexual differentiation in decapods.

The kuruma prawn *M. japonicus* (suborder Dendrobranchiata; family Penaeidae) is one of the most commercially important fishery species in Japan. In 2022, its production in Japan was approximately 214 tons from capture fisheries and 1,198 tons from aquaculture. In recent years, however, natural resources have been at risk of decline (Ministry of Agriculture, Forestry, and Fisheries of Japan, 2024). This species exhibits sexual dimorphism in growth, with females generally growing faster and larger than males (Coman et al., 2008). The development of all-female populations is a promising strategy for improving productivity in aquaculture. The full-length cDNA sequence of *M*. *japonicus* IAG, designated Maj-IAG, was reported (Banzai et al., 2011). Based on its expression site, AG is thought to be located at the distal end of the seminal vesicle (SV). Chemically synthesized Maj-IAG has been reported to suppress the expression of the major yolk protein gene, vitellogenin, in ovarian culture assays (Katayama et al., 2014), implying a possible role in diminishing female reproductive function. However, it remains unclear whether Maj-IAG is involved in male-intrinsic functions, such as the induction of male sexual differentiation and the maintenance of male sexual characteristics. Based on the findings in other decapod species, we hypothesized that Maj-IAG functions as an androgen in *M*. *japonicus* and orchestrates the development of both internal and external male traits. Therefore, the present study aimed to investigate the role of Maj-IAG in the development and maintenance of male secondary sexual characteristics using RNA interference (RNAi) in juvenile male *M*. *japonicus* after sexual differentiation.

## 2. Materials and methods

### 2.1. Animals

In this study, the binomial nomenclature of penaeid shrimp followed Pérez Farfante and Kensley (1997). Juvenile kuruma prawns *M*. *japonicus* were purchased from a local aquaculture company in Tokushima Prefecture, Japan, and transferred to the Mie Prefectural Fish Farming Center (Shima, Mie, Japan). Juveniles with a body length (BL) of 63.51 ± 3.6 mm and body weight (BW) of 2.79 ± 0.47 g (mean ± standard deviation), possessing the petasma, which is the male-specific external copulatory structure (Figure 1), but with underdeveloped testis, were used. They were divided into two groups, as described later, and stocked in two separate 200-L black circular tanks covered with light-shielding lids. Sand-filtered natural seawater (22.7–29.4℃) was continuously supplied to the tanks, and the prawns were fed a commercial diet (Goldprawn, Higashimaru Co., Ltd., Kagoshima, Japan) to satiation daily.

**Figure 1.**
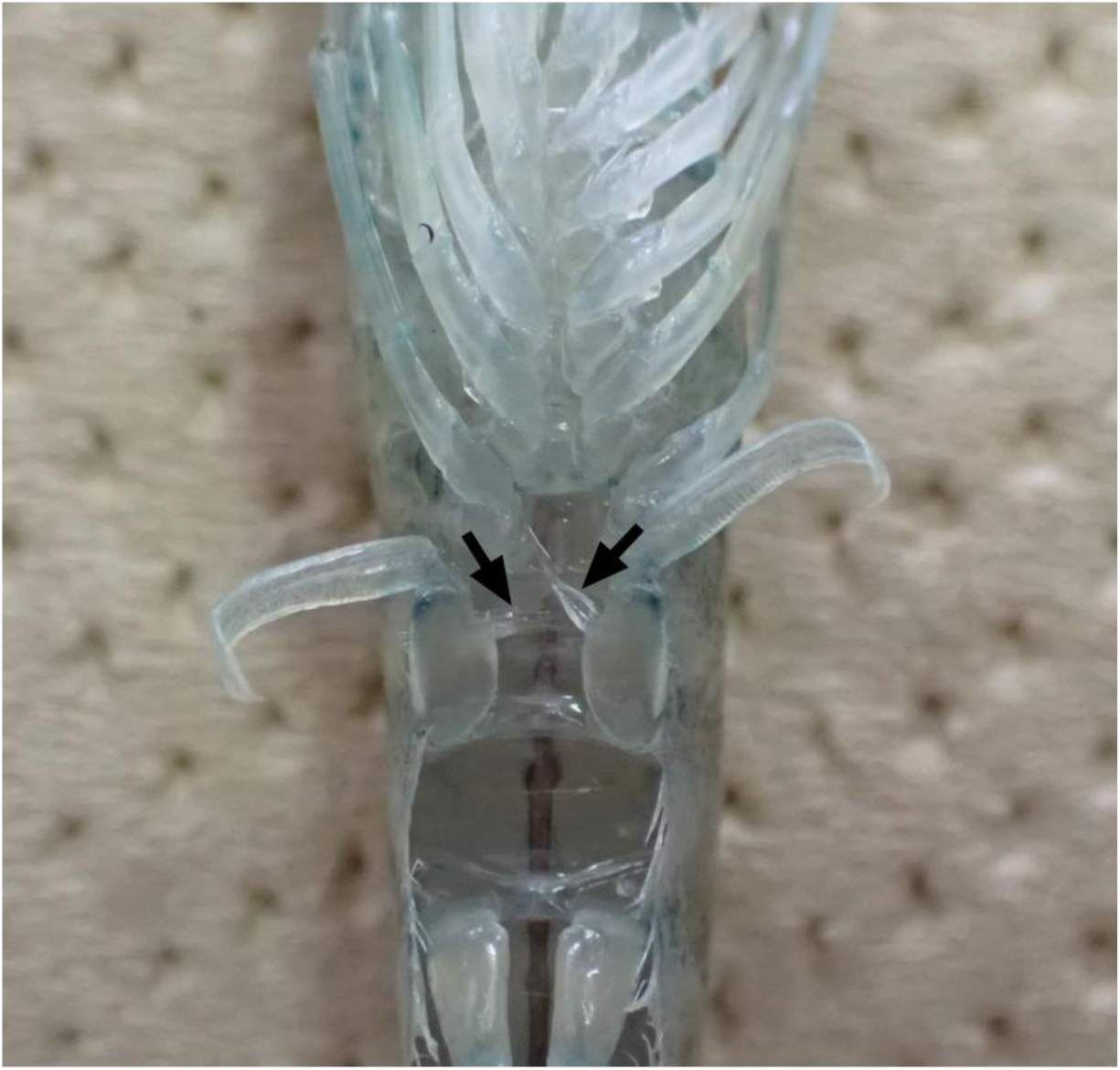
Ventral view of kuruma prawn (*Marsupenaeus japonicus*). Black arrows indicate the petasma, a male-specific copulatory structure formed on both the first pleopods.

### 2.2. Preparation of double-stranded RNA

Knockdown (KD) of *Maj-IAG* was achieved using RNAi with double-stranded RNA (dsRNA). dsRNA corresponding to the enhanced green fluorescent protein (EGFP) gene was used as a negative control. The sequence of *Maj-IAG* was retrieved from the NCBI database (accession number AB598415). Specific primer sets were then designed (Table 1) and templates for dsRNA synthesis were amplified by PCR using the primers. dsRNAs targeting *Maj-IAG* and *EGFP* were synthesized using the MEGAScript T7 Transcription Kit Plus (Invitrogen, Waltham, MA, USA) and purified using the Sepasol-RNA I Super G (Nacalai Tesque, Kyoto, Japan).

**Table 1.**
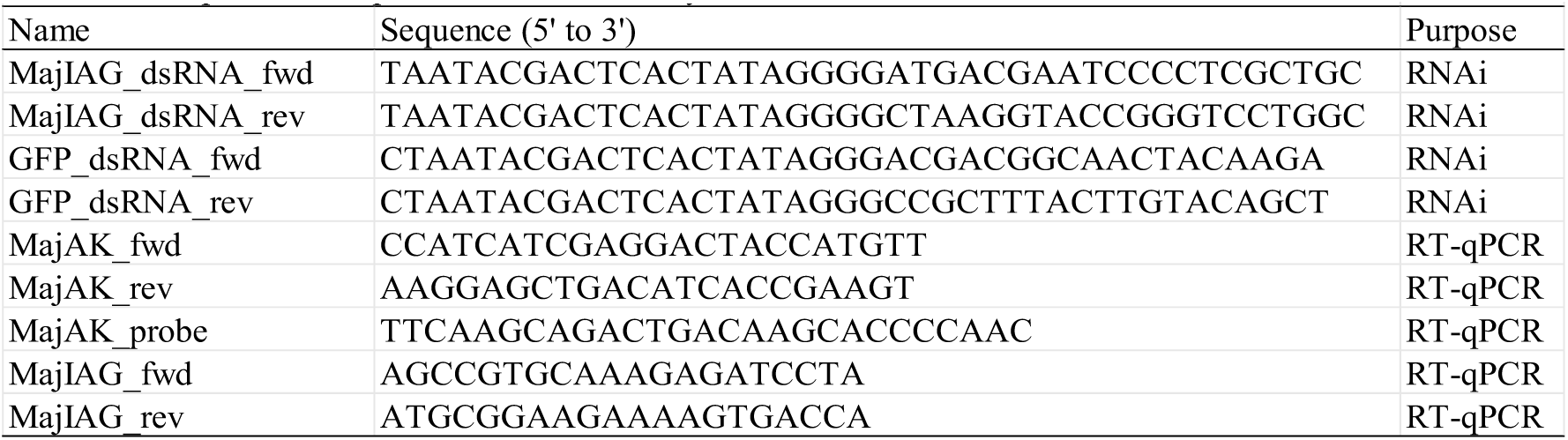
The primers and probe used in this study.

### 2.3. Gene knockdown and sampling schedule

Juvenile males were assigned to one of two groups: dsIAG and dsEGFP. dsRNA was injected into the body cavity of each prawn once every two weeks for a total of five times. Injections were performed using a micro-syringe equipped with an ultrafine needle (ITO CORPORATION, Shizuoka, Japan) under a stereomicroscope until week 6, and a disposable insulin syringe (Becton, Dickinson and Company, Franklin Lakes, NJ, USA) was used for injection at week 8. The dsRNA dose was gradually increased in accordance with prawn growth: 2, 3, 4, 5, and 6 μg/individual at the initial injection and at weeks 2, 4, 6, and 8, respectively.

At week 9 (one week after the final injection), the growth parameters BL and BW were measured. The reproductive organs were excised and divided into the testis, vas deferens (VD), and SV (Figure 2). A spermatophore is formed in the lumen of the SV. Testes were weighed to calculate the Testis Somatic Index (TSI; testis weight/BW × 100). Photographs of several VDs and SVs samples were captured using the iPod Touch 7th generation (Apple Inc., Cupertino, CA, USA) mounted on a stereomicroscope via i-NTER LENS (Micronet Corp., Saitama, Japan). SV samples were preserved in RNAlater solution (Thermo Fisher Scientific) and stored at −20℃ until RNA extraction. For histological analysis, the testis, VDs, and SVs were fixed in Bouin’s solution for 24 h and stored in 70% ethanol. Additionally, the first pleopods with petasmas were excised and fixed in Bouin’s solution for subsequent examination of the external sexual morphology.

**Figure 2.**
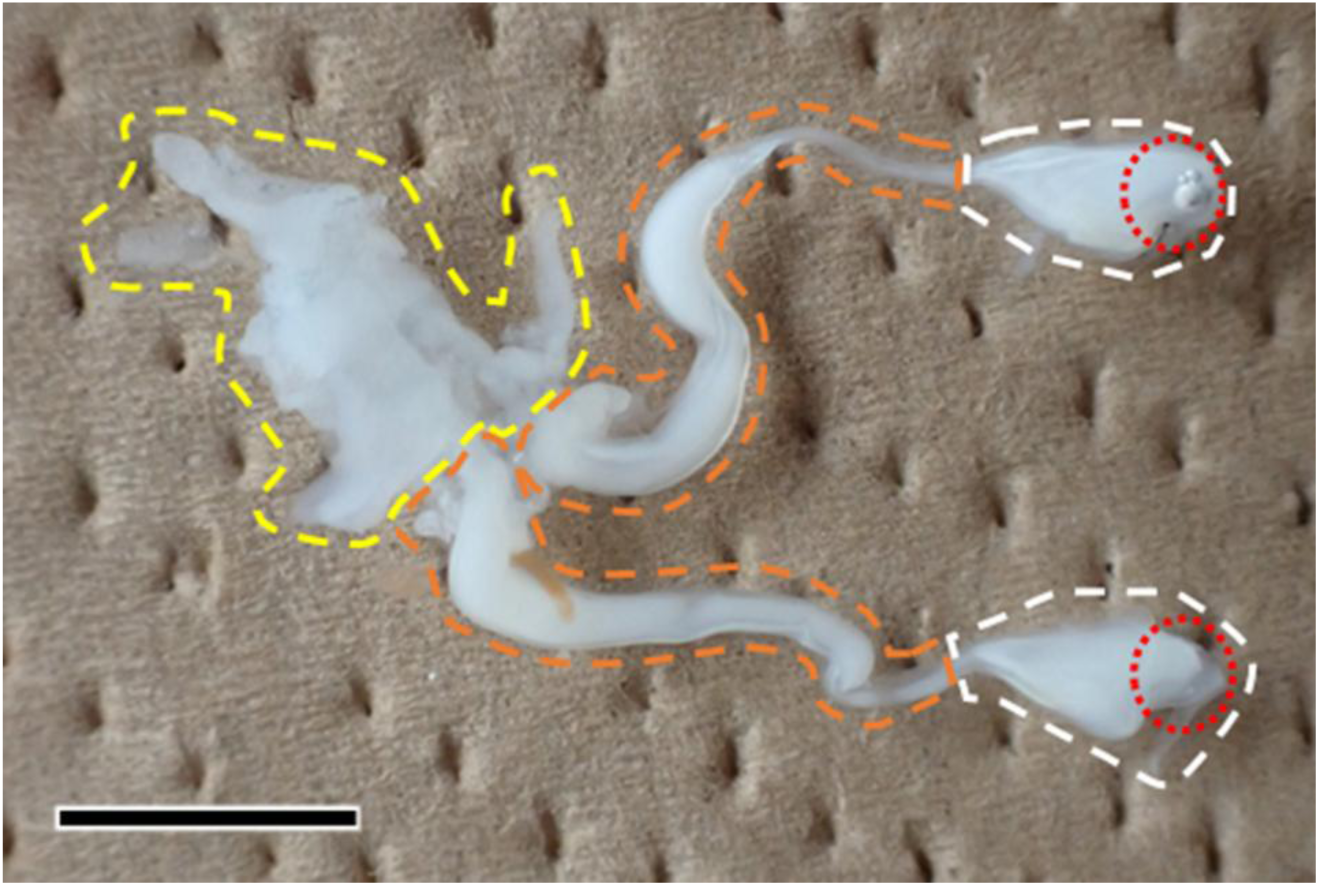
Photograph of male reproductive organs at week 9. This consists of three parts: the testis, vas deferens (VD), and seminal vesicle (SV), which are outlined by the yellow, orange, and white dashed lines, respectively. The androgenic gland (AG) is located at the distal end of the SV and is circled by red dots. A spermatophore is formed in the lumen of the SV. Scale bar: 50 µm (approximate because of the imaging conditions).

### 2.4. RNA extraction and quantitative RT-PCR

Total RNA was extracted using the NucleoSpin RNA isolation kit (Macherey–Nagel GmbH & Co. KG, Düren, Germany) according to the manufacturer’s instructions. Total RNA concentrations were measured using the NanoDrop Lite spectrophotometer (Thermo Fisher Scientific). The sequences of the primers and the TaqMan probe used for quantitative RT-PCR (RT-qPCR) are shown in Table 1. The *arginine kinase* (*Maj-AK*) was used as the internal reference gene followin Tsutsui et al. (2022). RT-qPCR was performed using the Luna Universal One-Step RT-qPCR kit (New England Biolabs, Ipswich, MA, USA) for *Maj-IAG* and the Luna Universal Probe One-Step RT-qPCR Kit (New England Biolabs) for *Maj-AK*. Each 20 µL reaction solution contained 4 ng of total RNA and 400 nM of each primer; in the case of *Maj-AK*, 200 nM of TaqMan probe was also included.

Amplification and real-time fluorescence detection were performed using the 7300 Real-Time PCR System (Applied Biosystems, Foster City, CA, USA) under the following thermal cycling conditions: reverse transcription at 55℃ for 10 min, initial denaturation at 95℃ for 1 min, followed by 45 cycles of 95℃ for 15 s and 60℃ for 60 s. All reactions were performed in duplicate. Relative gene expression levels were calculated using the 2^−ΔΔCt^ method, with expression levels normalized to *Maj-AK*.

### 2.5. Measurement of tissue length and area

Photographs of the fixed petasmas were captured using the imaging system described above. Petasma length was measured using Fiji/ImageJ software version 1.54f (Schindelin et al., 2012; Schneider et al., 2012). In addition, the length of the VD and the areas of the VD and SV were analyzed. For each prawn, the left and right sides of the paired reproductive organs were measured, and the average value was used as the data for each individual. Measurements of length and area were normalized to BL and (BL)^2^, respectively, to account for individual size differences.

### 2.6. Histology of male reproductive organs

The fixed tissues were dehydrated using an ethanol series and embedded in paraffin. Serial sections (5–8 µm thickness) were prepared using the RM2125RT microtome (Leica, Bensheim, Germany) and stained with hematoxylin and eosin. Tissue sections were observed under the Axio Imager.A1 light microscope (Carl Zeiss, Oberkochen, Germany), and digital images were captured using the Axiocam 208 color camera (Carl Zeiss) mounted on the microscope.

### 2.7. Statistics

Statistical analysis was performed using Welch’s t-test in R (version 4.1.2; R Core Team, 2023) with RStudio (version 2021.09.2+382; Posit team, 2023) to compare all data between the dsIAG and dsEGFP groups. Statistical significance was set at *p* < 0.05. All data were used for boxplot drawing without outlier calculations. Statistical analysis of RT-qPCR data was performed based on ΔCt values.

## 3. Results

### 3.1. Knockdown efficiency of *Maj-IAG* and its effects on somatic growth

To assess KD efficiency, the expression levels of *Maj-IAG* in the SV with AG at week 9 were measured using RT-qPCR. Relative *Maj-IAG* expression in the dsIAG group was significantly reduced by approximately 65% compared to that in the dsEGFP control group (Figure 3). All prawns showed substantial growth from week 0 (BL: 63.51 ± 3.6 mm, BW: 2.79 ± 0.47 g). However, no statistically significant differences in BL were observed between the dsIAG group (92.89±3.61 mm) and the dsEGFP group (95.34±4.26 mm) at week 9 (Figure 4A). Similarly, BW did not differ significantly between the two groups at week 9 (9.67±1.17 g in dsIAG vs. 10.04±1.32 g in dsEGFP) (Figure 4B).

**Figure 3.**
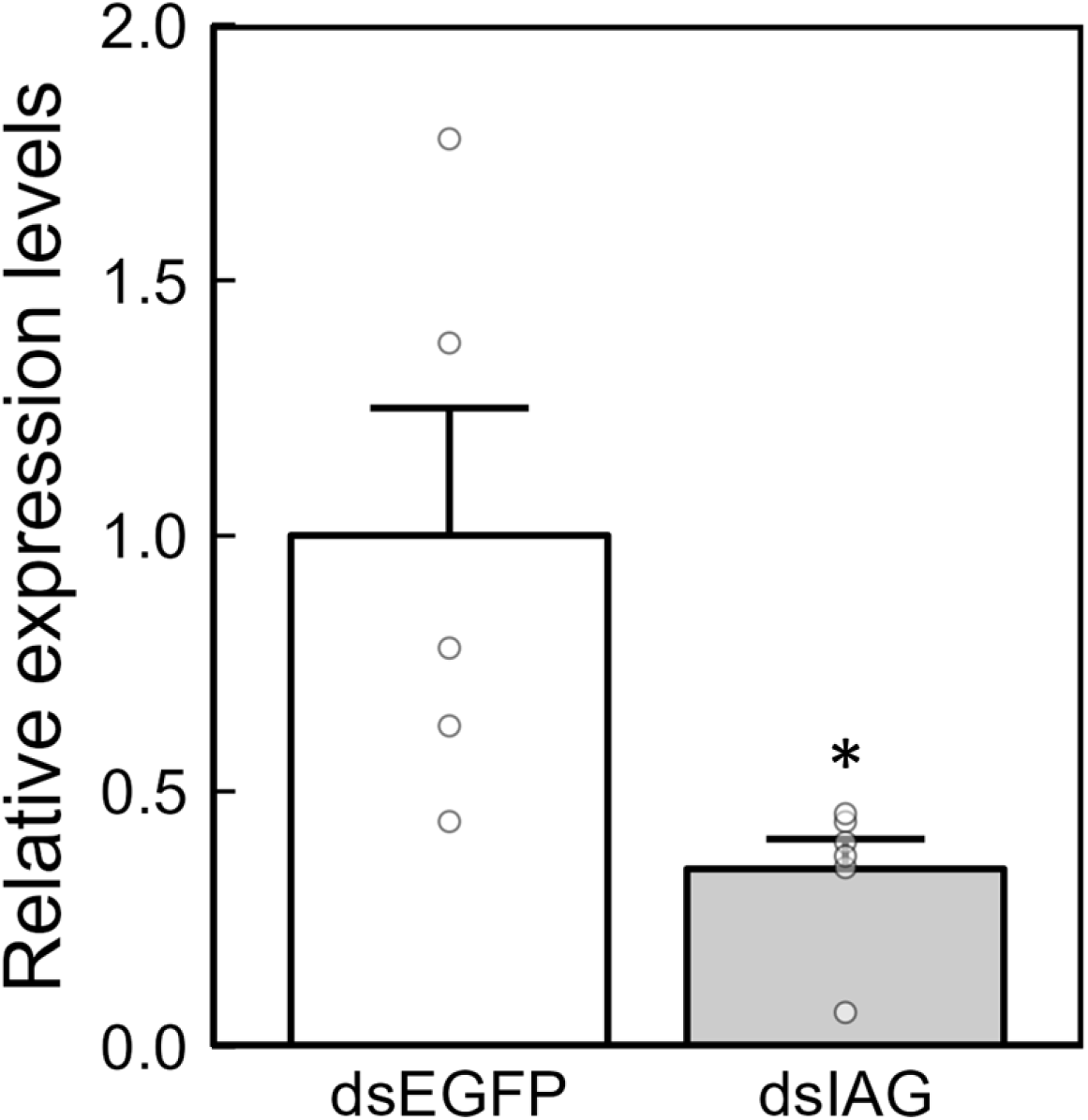
The knockdown efficiency of *Maj-IAG* following dsIAG administration. Relative expression levels of *Maj-IAG* in the SV, including the AG, were normalized to those of *Maj-AK* and are represented as mean + SEM (dsEGFP: *n* = 5, dsIAG: *n* = 6). Dot plots show individual values. An asterisk indicates a significant difference between the dsIAG and dsEGFP groups, as determined by Welch’s t-test (\**P* < 0.05).

**Figure 4.**
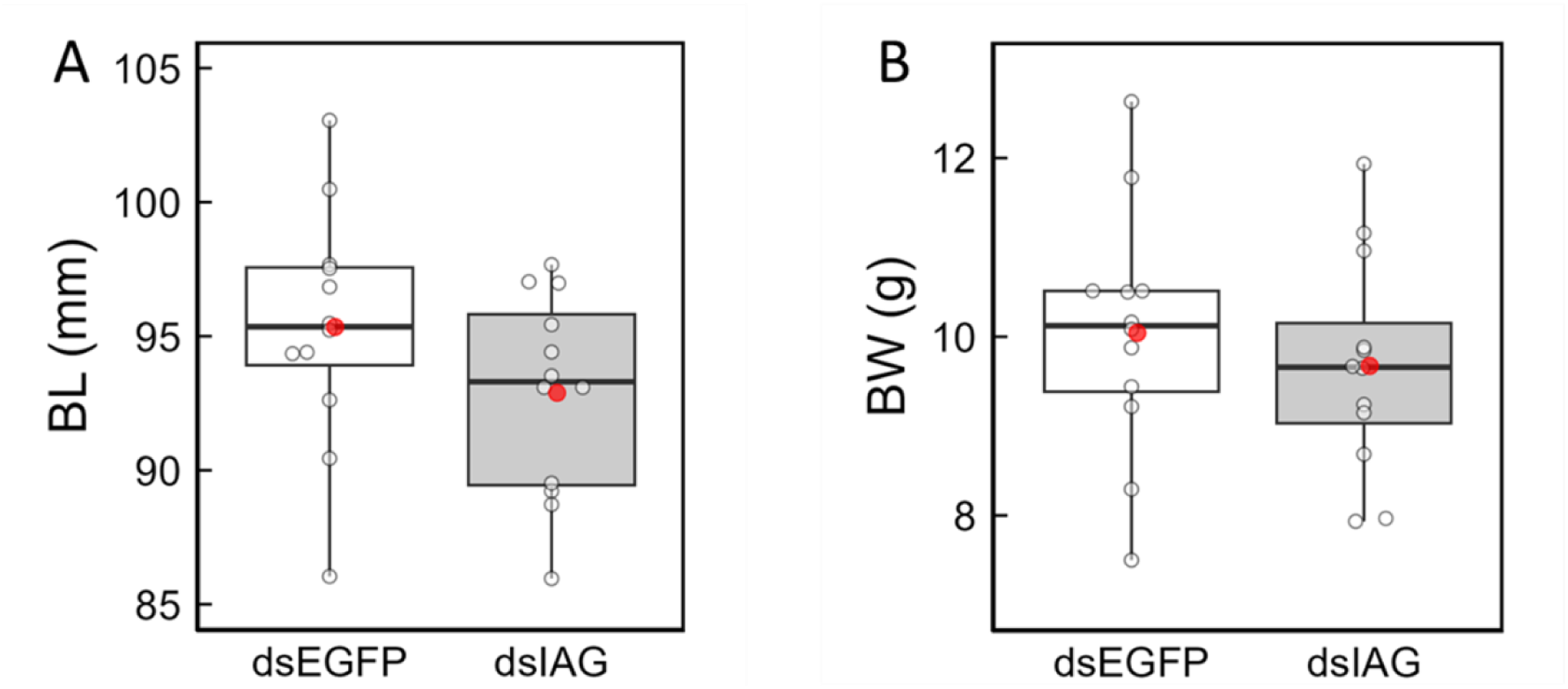
Effects of *Maj-IAG* knockdown on somatic growth. Boxplots show BL (A) and BW (B), with individual values overlaid as dot plots and mean values indicated by red dots (*n* = 12 per group). There were no significant differences in the BL and BW between the dsIAG and dsEGFP groups by Welch’s t-test (*P* > 0.05).

### 3.2. Effects of *Maj-IAG* knockdown on the development of external masculine characteristics

The development of petasma, a primary external male characteristic, was examined to assess whether *Maj-IAG* KD influences external sexual morphology. Macroscopic observation at week 9 revealed no apparent petasma deformities in either the dsIAG or dsEGFP groups. Therefore, a quantitative analysis of petasma length was conducted using photographs (Figure 5A). Because *Maj-IAG* KD did not affect the overall BL (Figure 4A), petasma length was normalized to the BL of the individual. The resulting petasma length: BL ratio was significantly lower in the dsIAG group than in the dsEGFP group (Figure 5B).

**Figure 5.**
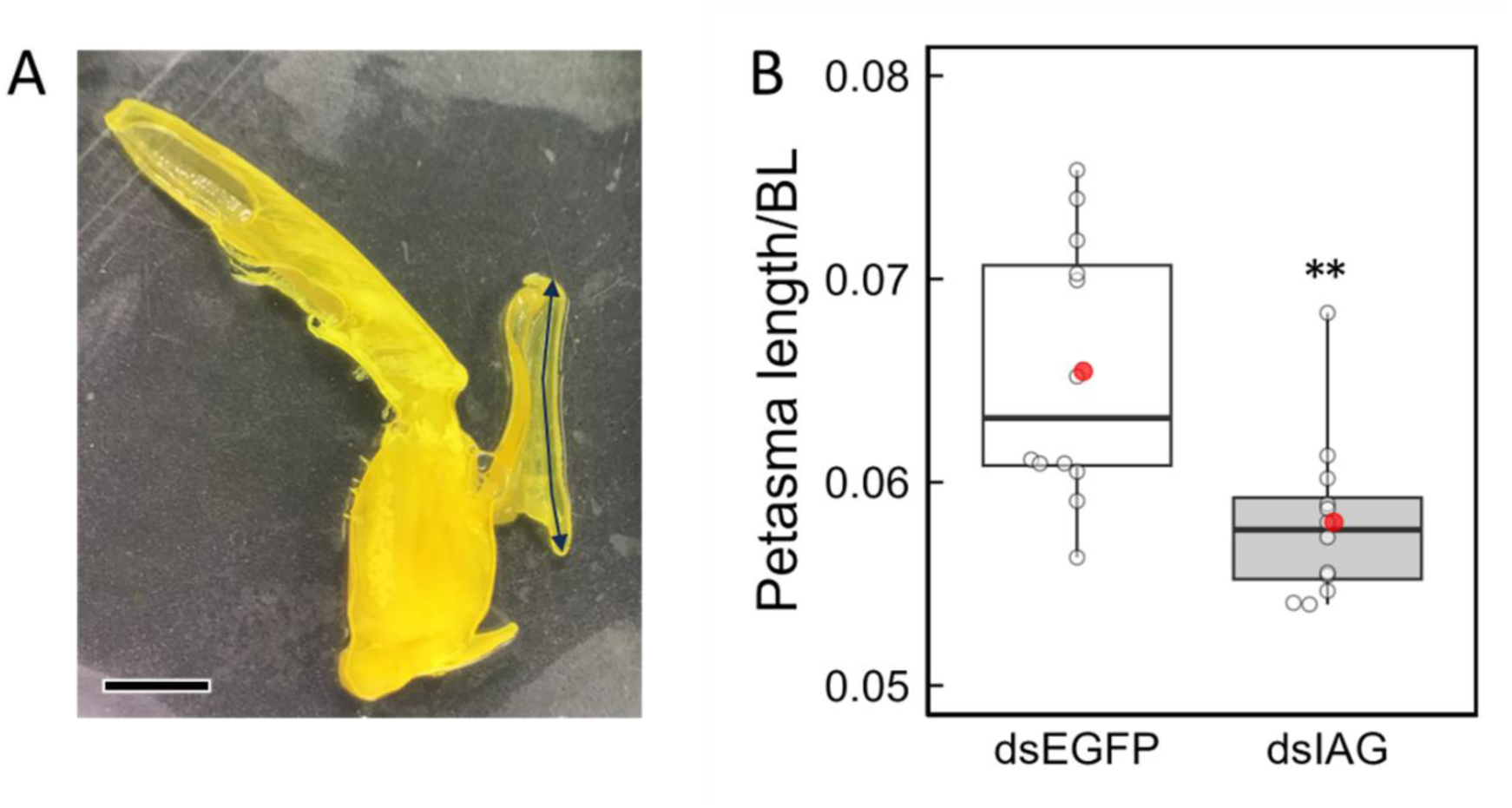
The development of petasma under *Maj-IAG* knockdown. The photograph shows a fixed right first pleopods (A). The petasma length is indicated by a double-headed arrow. The scale bar represents 2.5 mm. The boxplots represent petasma length normalized by BL. Individual values are shown as dot plots, and the means (*n* = 12 per group) are indicated by red dots (B). Asterisks indicate significant differences between the dsIAG and dsEGFP groups, as determined by Welch’s t-test (\*\**P* < 0.01).

### 3.3. Effects of *Maj-IAG* knockdown on the development of internal male reproductive organs

Morphological and quantitative analyses were conducted to evaluate the effects of *Maj-IAG* KD on the internal development of male reproductive structures. Among the three major parts of the male reproductive system, namely the testis, VD, and SV, only the testis had a measurable weight at week 9. However, the TSI did not differ significantly between the dsIAG and dsEGFP groups (Figure 6). For VD and SV, whose weights could not be measured, a developmental assessment was conducted by analyzing digital images captured under a stereomicroscope (Figure 7A, B). The length of the VD and areas of both the VD and SV were measured and normalized to each BL or (BL)^2^ of the individual. All these parameters (VD length/BL, VD area/(BL)^2^, and SV area/(BL)^2^) were significantly lower in the dsIAG group compared to the dsEGFP group (Figure 7C–E).

**Figure 6.**
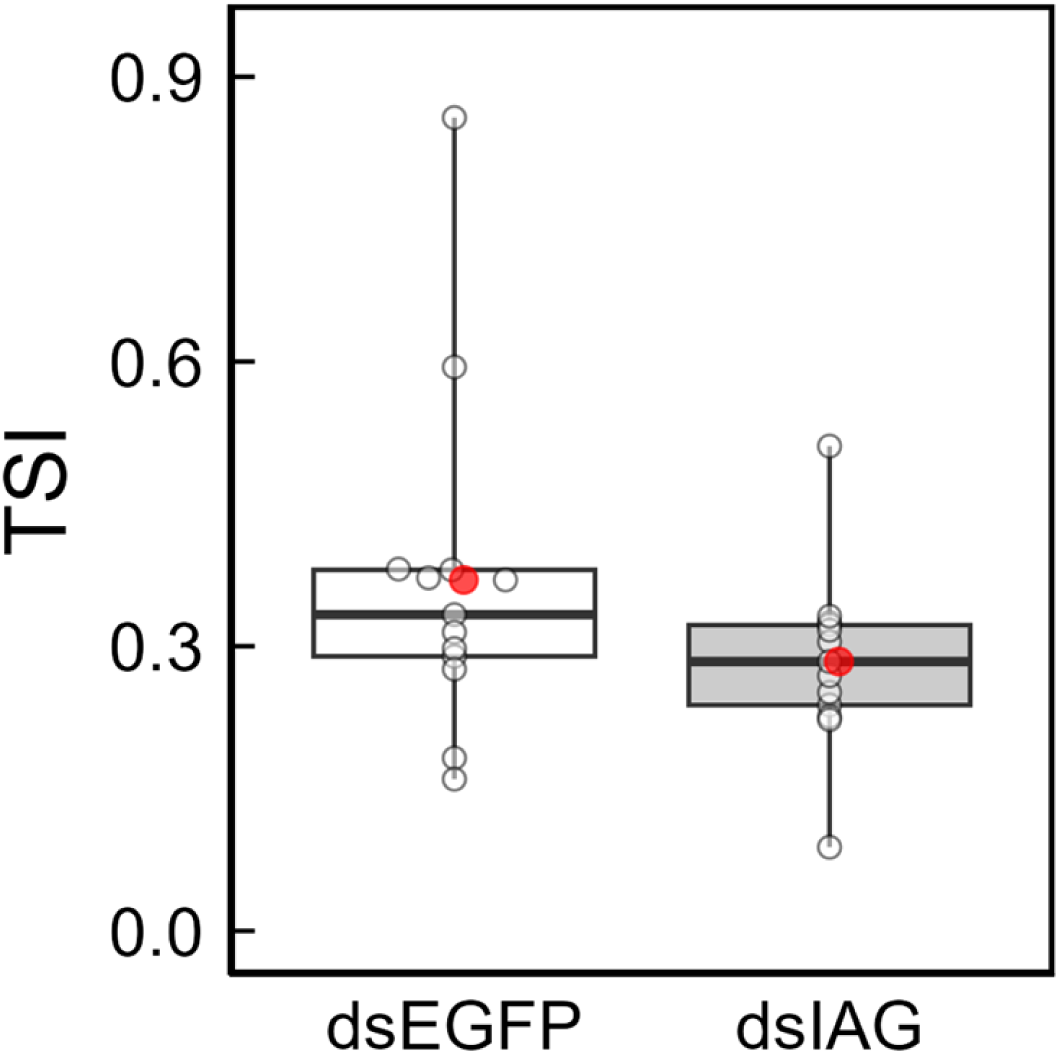
Effects of *Maj-IAG* knockdown on Testis Somatic Index (TSI). The boxplots show individual values as dot plots, with the means (*n* = 12 per group) indicated by the red dots. There were no significant differences in TSI between the dsIAG and dsEGFP groups by Welch’s t-test (*P* > 0.05).

**Figure 7.**
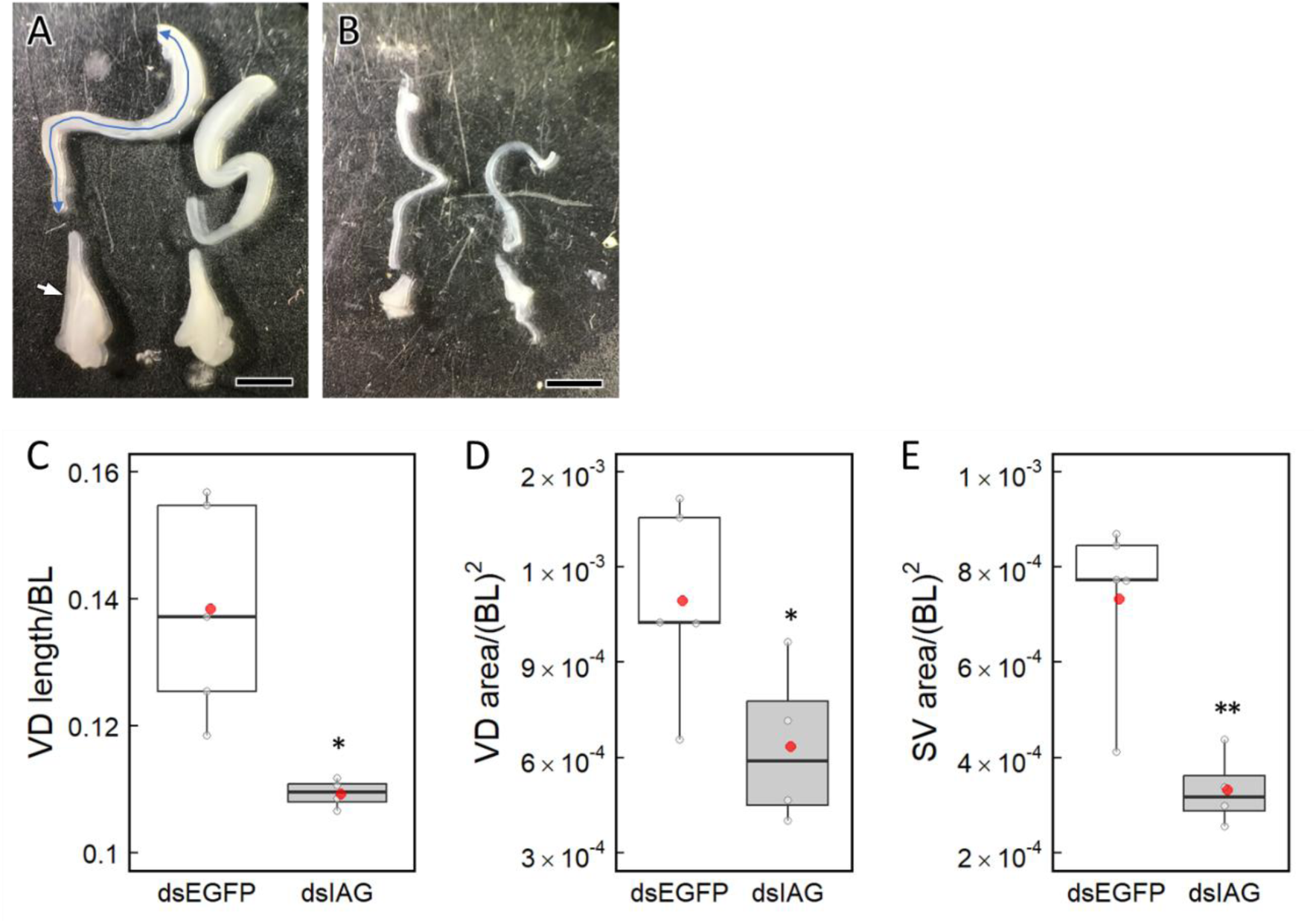
Effects of *Maj-IAG* knockdown on the development of male reproductive organs. (A) and (B) show representative photographs of the reproductive organs, the VD and SV, in the dsEGFP and dsIAG groups, respectively. Scale bars represent 2.5 mm. The double-headed arrow indicates the VD length as defined in this study, and the white arrow indicates the SV. The boxplots show individual values normalized to BL or (BL)^2^ as dot plots (C-E). Red dots show mean values (dsEGFP: *n* = 5, dsIAG: *n* = 4). Asterisks indicate significant differences between the dsEGFP and dsIAG groups by Welch’s t-test (\**P* < 0.05, \*\**P* < 0.01).

### 3.4. Histological analysis of the male reproductive organ

Histological analysis of the testis was performed to examine whether *Maj-IAG* KD affected spermatogenesis. Spermatogonia, primary spermatocytes, and spermatids undergoing transformation into spermatozoa were observed in both dsEGFP and dsIAG groups at week 9 (Figure 8A, B, and S1A–B). Although the seminiferous tubule area of the testes was not significantly different between the two groups, it was larger than that observed at week 0 (Figure 8A, B, and S1C). Compared to the initial group, germ cell development progressed during the 9-week rearing period. However, no histological differences were observed in the testes between the two groups at week 9. In VD, although the precise sectioning sites may not have been fully consistent among individuals, the duct diameter appeared to differ between the dsIAG and dsEGFP groups (Figure 8C, D). No differences were observed in the internal tissue structure, and spermatozoa were present in both groups (Figure 8E–F). In SV, control prawns showed spermatophore formation and the presence of spermatozoa (Figure S1D and F). In contrast, no structure corresponding to the spermatophore was observed in *Maj-IAG* KD prawns, although this may have resulted from slight differences in the sectioning sites (Figure S1E).

**Figure 8.**
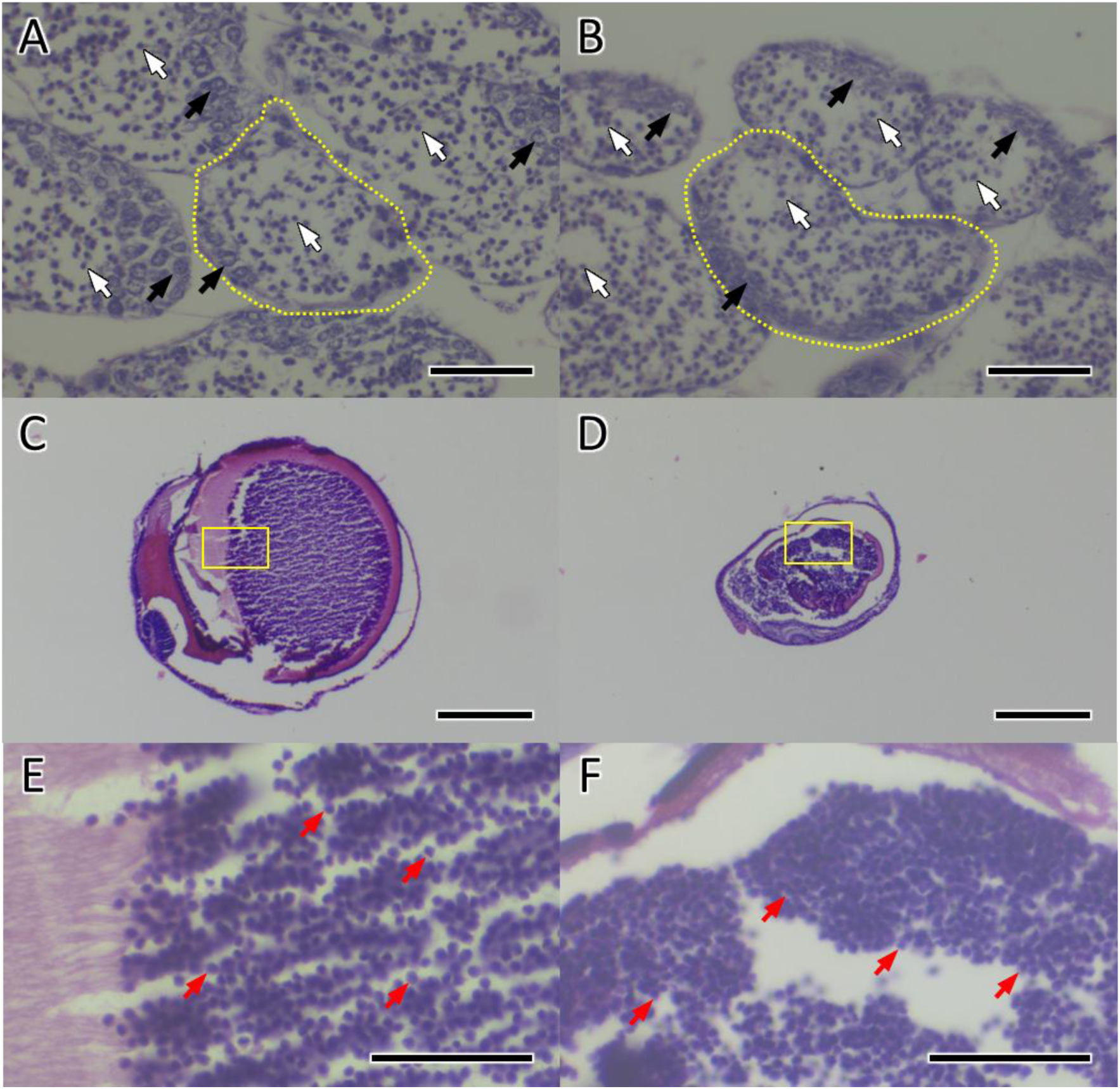
Histological analysis of the testes and VDs at week 9. Representative cross-sections of the testis (A, B) and VD (C, D) with enlarged views of spermatozoa in the VD (E, F) are shown. Yellow squares in (C) and (D) indicate the magnified regions in (E) and (F), respectively. Panels (A), (C), and (E) correspond to the dsEGFP group, whereas panels (B), (D), and (F) correspond to the dsIAG group. Black arrows indicate spermatogonia, white arrows indicate spermatids, and red arrows indicate spermatozoa. A typical cross-section of a testicular seminiferous tubule is outlined by yellow dotted lines. Scale bars: 50 µm in (A, B, E, F); 250 µm in (C, D).

## 4. Discussion

The results of the present study demonstrate that Maj-IAG plays a physiological role in the development of male secondary sexual characteristics in the juvenile kuruma prawn *M*. *japonicus* (suborder Dendrobranchiata; family Penaeidae). Quantitative assessment of the developmental degree of each reproductive organ revealed that *Maj-IAG* KD suppressed the development of petasma, VD, and SV in juvenile males. To the best of our knowledge, this is the first direct evidence of a specific function of IAG in males of this species. Although the methods used to alter endogenous IAG signaling differed, with IAG gene silencing in *M*. *japonicus* in our study and AG ablation in *L*. *vannamei* (suborder Dendrobranchiata; family Penaeidae) in the study by Alfaro-Montoya et al. (2016), inhibition of petasma development was similarly observed in both penaeid species. In the peppermint shrimp *Lysmata vittata* (suborder Pleocyemata; infraorder Caridea), *IAG* silencing similarly inhibited the development of appendix masculina (Liu et al., 2021). In the red claw crayfish *Cherax quadricarinatus* (suborder Pleocyemata; infraorder Astacidea) and *M*. *rosenbergii* (suborder Pleocyemata; infraorder Caridea), AG ablation induced not only the inhibition of appendix masculina development but also VD abnormalities in *C*. *quadricarinatus* and *M*.*rosenbergii* (Nagamine et al., 1980; Tropea et al., 2011). Taken together with our findings, these results suggest that IAG plays a conserved role in the development of male reproductive organs in decapod crustaceans.

It is thought that IAG is involved in growth regulation in relation to sexual size dimorphism in decapod crustaceans. In *M*. *rosenbergii*, which displays a sexually dimorphic growth pattern in larger males (Sagi et al., 1986), growth inhibition following AG ablation or *IAG* silencing has been observed (Sagi and Cohen, 1990; Ventura et al., 2012, 2009). In *C*. *quadricarinatus*, another species in which males exhibit a significant growth advantage, AG implantation into females increased their growth rate to levels comparable to that of males (Manor et al., 2004). Moreover, in *L*. *vannamei*, a species with female-biased growth dimorphism (Moss and Moss, 2006), growth inhibition has been observed in males following AG ablation (Alfaro-Montoya et al., 2016). Li et al. (2012) noted that both IAG and insulin-like growth factor belong to the insulin superfamily, and proposed that IAG may also promote growth. Simultaneously, they indicated that as a male-specific factor, IAG may not be essential for growth (Li et al., 2012). In the present study, *Maj-IAG* KD had no significant effect on somatic growth, suggesting that Maj-IAG is not involved in the growth regulation of male *M*. *japonicus*. Similarly, in male *C*. *quadricarinatus*, no growth effect was observed following AG ablation (Tropea et al., 2011). The extent to which IAG signaling contributes to the regulation of sexual size dimorphism may vary across species, potentially depending on which sex exhibits a growth advantage, indicating the need for further detailed investigation in each species. Regarding germ cell differentiation in the testis, spermatids undergoing transformation into spermatozoa were the most advanced stage observed in both the dsIAG and dsEGFP groups, and the progression of differentiation from spermatogonia to spermatozoa during the experimental period was evident. The absence of marked differences in spermatogenic stages by histology, along with comparable testicular development indicated by similar TSI values in both the dsIAG and dsEGFP groups, suggests that Maj-IAG may not be involved in the development, maintenance, or function of the testis. This is supported by the fact that *Maj-IAG* KD in juveniles for 9 weeks resulted in no obvious abnormalities, including testicular degeneration or abnormal arrest of germ cell differentiation, both of which are indicative of male reproductive dysfunction. Moreover, a previous study on the same species suggested that testis differentiation and development are driven by genetic sex determination, without the involvement of young AG (Nakamura et al., 1992). In contrast, the KD of a putative IAG receptor in the Chinese shrimp *Fenneropenaeus chinensis* (suborder Dendrobranchiata; family Penaeidae) led to the arrest of germ cell development at the secondary spermatocyte stage (Guo et al., 2019). *IAG* silencing in *P*. *clarkii* suppressed spermatogenesis (Ge et al., 2020). AG ablation in juvenile *L*. *vannamei* induced testicular degeneration and decreased TSI (Alfaro-Montoya et al., 2016). These findings highlight the need for comparative studies across diverse decapod species to clarify the role of the IAG signaling pathway in spermatogenesis and testicular development. In the present study, histologically normal spermatozoa were found to be present in VDs; that is, the sperm transport pathway from the testis to the SVs was unaffected, even under *Maj-IAG* KD. However, the structural development of VDs and SVs was impaired. These results suggest that IAG signaling in *M. japonicus* may be more closely involved in the morphological maturation of these reproductive ducts than during spermatogenesis.

The results of the present study suggest that IAG plays an important role in the expression of secondary sexual characteristics in *M*. *japonicus*, whereas testicular development and maturation after sex differentiation may be controlled by a robust mechanism independent of IAG signaling. In *M*. *rosenbergii*, IAG silencing has been reported to successfully induce sex reversal and subsequent gametogenesis (Ventura et al., 2012). In the juvenile Chinese mitten crab *Eriocheir sinensis* (suborder Pleocyemata; infraorder Brachyura), feminization of the external morphology was observed after *IAG* KD (Fu et al., 2020). In certain crayfish species (suborder Pleocyemata; infraorder Astacidea), including the marbled crayfish *Procambarus virginalis* and *P*. *clarkii*, IAG has been shown to suppress oocyte maturation (Katayama et al., 2022; Taketomi and Nishikawa, 1996). Furthermore, a more recent study demonstrated that *IAG* knockout using the CRISPR/Cas9 system in the male ridgetail white prawn *Exopalaemon carinicauda* (suborder Pleocyemata; infraorder Caridea) successfully induced sex reversal and development of phenotypic female characteristics (Miao et al., 2023). In contrast, in juvenile *L*. *vannamei*, AG ablation has been reported to affect the development of sexual characteristics, such as testicular degeneration and decreased TSI, but sex reversal was not induced (Alfaro-Montoya et al., 2016). Similarly, in females implanted with AG 34–37 days after metamorphosis into the post-larval stage, some individuals exhibited petasma formation, whereas abnormal oocyte development, oocyte regression, or functional masculinization were not observed (Vega-Alpízar et al., 2017). It is plausible that IAG functions as a hormonal regulator of sex differentiation in decapod crustaceans; however, its effects have not been consistently observed across species within this taxon, even when differences in experimental procedures are considered. Based on the findings of this study and previous reports, the regulatory mechanisms of gametogenesis in the suborder Dendrobranchiata, including *M*. *japonicus* and *L*. *vannamei,* may be more robust and less susceptible to IAG manipulation. To date, there have been no reports of complete loss of male sexual function or successful sex reversal in Dendrobranchiata species, suggesting relatively low sexual plasticity in their reproductive organs compared to members of Pleocyemata. This idea is supported by Alfaro-Montoya et al. (2019), who suggested that sexual plasticity varies among taxonomic groups. Through phylogenetic analysis using mitochondrial genomes, researchers have estimated that Pleocyemata and Dendrobranchiata diverged approximately 423 million years ago (Porter et al., 2005) and that differences in sexual plasticity may have emerged during the course of their speciation. Further IAG-focused studies across a broad range of species in both Dendrobranchiata and Pleocyemata are required to better understand the mechanisms underlying sexual plasticity in reproductive organs.

In this study, *Maj-IAG* expression levels were suppressed by an average of 65%; however, on an individual basis, some individuals in the dsEGFP group exhibited low *Maj-IAG* expression levels comparable to those in the dsIAG group. This variability may reflect differences in the developmental stage of the AG, which was still developing in juveniles, leading to natural variation in *Maj-IAG* expression during the experimental period. Despite this variation, significant differences in the development of petasma, VD, and SV were observed between the two groups, highlighting the physiological impact of *Maj-IAG* KD. In contrast, the KD efficiency of *Maj-IAG* achieved in this study may not be sufficient to reveal the full role of IAG in testicular development and spermatogenesis. The absence of obvious abnormalities, such as testicular degeneration or abnormal arrest of germ cell differentiation, suggests that the residual IAG signal in the dsIAG group may have been adequate to sustain normal spermatogenesis and testicular development. To clarify whether IAG signaling plays an essential role in spermatogenesis and testis development in *M*. *japonicus*, a more efficient knockdown approach or complete gene knockout may be necessary. In addition, identification of the IAG receptor and subsequent knockdown or knockout of the corresponding gene may be required.

## 5. Conclusion

In this study, the physiological role of *Maj-IAG* in the development of male secondary sexual characteristics in the juvenile kuruma prawn *M*. *japonicus* was determined using RNAi. Specifically, *Maj-IAG* was found to promote the development of petasma, VD, and SV in juveniles, providing the first direct evidence of its functional significance in the males of this species. The role of IAG in the regulation of male secondary sexual characteristics may be conserved among decapod crustaceans. In contrast, the phenotypic differences resulting from IAG manipulation across species suggest that sexual plasticity may vary not only among broader taxonomic groups but also among closely related species. This study provides fundamental insights into the potential development of sex-control technology in *M*. *japonicus*, with broader implications for penaeid shrimps, for which functional sex reversal has not yet been achieved.

## CRediT authorship contribution statement

**Takehiro Furukawa:** Conceptualization, Writing – original draft, Writing – review & editing, Methodology, Investigation, Formal analysis, Data curation, Visualization, Funding acquisition. **Fumihiro Yamane:** Writing – review & editing, Investigation, Resources**. Rino Takeuchi:** Writing – review & editing, Investigation. **Tsuyoshi Ohira:** Writing – review & editing, Resources, Methodology. **Kenji Toyota:** Writing – review & editing, Methodology. **Taeko Miyazaki:** Writing – review & editing, Resources, Methodology. **Naoaki Tsutsui:** Writing – review & editing, Supervision, Project administration, Funding acquisition.

## Acknowledgments

This research was helped by Mie Prefectural Fish Farming Center for their kind support of daily life during the experiment. Valuable advice on germline cell staging was kindly provided by Dr. Takuji Okumura (The Japan Fisheries Research and Education Agency). The study was supported by a Grant-in-Aid for Scientific Research [grant number 22K05831] from the Ministry of Education, Culture, Sport, Science, and Technology (MEXT) of Japan and JST SPRING [grant number JPMJSP2137].

## Data availability

All data generated or analyzed during this study are included in this published article or in the supplementary information files.

## References

Alfaro-Montoya, J., Braga, A., Umaña-Castro, R., 2019. Research frontiers in penaeid shrimp reproduction: Future trends to improve commercial production. Aquac. 503, 70–87. 10.1016/j.aquaculture.2018.12.068

Alfaro-Montoya, J., Hernández-Noguera, L., Vega-Alpízar, L., Umaña-Castro, R., 2016. Effects of androgenic gland ablation on growth, sexual characters and spermatogenesis of the white shrimp, *Litopenaeus vannamei* (Decapoda: Penaeidae) males. Aquac. Res. 47, 2768–2777. 10.1111/are.12727

Banzai, K., Ishizaka, N., Asahina, K., Suitoh, K., Izumi, S., Ohira, T., 2011. Molecular cloning of a cDNA encoding insulin-like androgenic gland factor from the kuruma prawn *Marsupenaeus japonicus* and analysis of its expression. Fish. Sci. 77, 329–335. 10.1007/s12562-011-0337-8

Coman, F.E., Sellars, M.J., Norris, B.J., Coman, G.J., Preston, N.P., 2008. The effects of triploidy on *Penaeus* (*Marsupenaeus*) *japonicus* (Bate) survival, growth and gender when compared to diploid siblings. Aquac. 276, 50–59. 10.1016/j.aquaculture.2008.01.031

Fu, C., Li, F., Wang, L., Wu, F., Wang, J., Fan, X., Liu, T., 2020. Molecular characteristics and abundance of insulin-like androgenic gland hormone and effects of RNA interference in *Eriocheir sinensis*. Anim. Reprod. Sci. 215, 106332. 10.1016/j.anireprosci.2020.106332

Ge, H.L., Tan, K., Shi, L.L., Sun, R., Wang, W.M., Li, Y.H., 2020. Comparison of effects of dsRNA and siRNA RNA interference on insulin-like androgenic gland gene (IAG) in red swamp crayfish *Procambarus clarkii*. Gene 752, 144783. 10.1016/j.gene.2020.144783

Guo, Q., Li, S., Lv, X., Xiang, J., Manor, R., Sagi, A., Li, F., 2019. Sex-biased CHHs and their putative receptor regulate the expression of IAG gene in the shrimp *Litopenaeus vannamei*. Front. Physiol. 10, 1–14. 10.3389/fphys.2019.01525

Katayama, H., Kubota, N., Hojo, H., Okada, A., Kotaka, S., Tsutsui, N., Ohira, T., 2014. Direct evidence for the function of crustacean insulin-like androgenic gland factor (IAG): Total chemical synthesis of IAG. Bioorganic and Medicinal Chemistry 22, 5783–5789. 10.1016/j.bmc.2014.09.031

Katayama, H., Toyota, K., Tanaka, H., Ohira, T., 2022. Chemical synthesis and functional evaluation of the crayfish insulin-like androgenic gland factor. Bioorg. Chem. 122, 105738. 10.1016/j.bioorg.2022.105738

Kinami, R., Ineno, T., 2025. First evidence of WW superfemale for hybrid sturgeon, the bester (*Huso huso* × *Acipenser ruthenus*). Aquac. 596, 741808. 10.1016/j.aquaculture.2024.741808

Levy, T., Aflalo, E.D., Sagi, A., 2018. Sex control in cultured decapod crustaceans, in: Sex Control in Aquaculture. John Wiley & Sons, Ltd, pp. 689–704. 10.1002/9781119127291.ch35

Levy, T., Sagi, A., 2020. The “IAG-Switch”—A Key Controlling Element in Decapod Crustacean Sex Differentiation. Front. Endocrinol. 11, 1–15. 10.3389/fendo.2020.00651

Li, S., Li, F., Sun, Z., Xiang, J., 2012. Two spliced variants of insulin-like androgenic gland hormone gene in the Chinese shrimp, *Fenneropenaeus chinensis*. Gen. Comp. Endocrinol. 177, 246–255. 10.1016/j.ygcen.2012.04.010

Liu, F., Shi, W., Ye, H., Zeng, C., Zhu, Z., 2021. Insulin-like androgenic gland hormone 1 (IAG1) regulates sexual differentiation in a hermaphrodite shrimp through feedback to neuroendocrine factors. Gen. Comp. Endocrinol. 303, 113706. 10.1016/j.ygcen.2020.113706

Mair, G.C., Abucay, J.S., Abella, T.A., Beardmore, J.A., Skibinski, D., 1997. Genetic manipulation of sex ratio for the large-scale production of all-male tilapia *Oreochromis niloticus*. Can. J. Fish. Aquat. Sci. 54, 396–404. 10.1139/f96-282

Mair, G.C., Abucay, J.S., Beardmore, J.A., Skibinski, D.O.F., 1995. Growth performance trials of genetically male tilapia (GMT) derived from YY-males in *Oreochromis niloticus* L.: On station comparisons with mixed sex and sex reversed male populations. Aquaculture, Genetics in Aquaculture V Proceedings of the Fifth International Symposium on Genetics in Aquaculture 137, 313–323. 10.1016/0044-8486(95)01110-2

Malecha, S.R., Nevin, P.A., Ha, P., Barck, L.E., Lamadrid-Rose, Y., Masuno, S., Hedgecock, D., 1992. Sex-ratios and sex-determination in progeny from crosses of surgically sex-reversed freshwater prawns, Macrobrachium rosenbergii. Aquac. 105, 201–218. 10.1016/0044-8486(92)90087-2

Manor, R., Aflalo, Eliahu D., Segall, Carmen, Weil, Simy, Azulay, Dudu, Ventura, Tomer, and Sagi, A., 2004. Androgenic gland implantation promotes growth and inhibits vitellogenesis in *Cherax quadricarinatus* females held in individual compartments. Invertebr. Reprod. Dev. 45, 151–159. 10.1080/07924259.2004.9652584

Matsunaga, T., Ieda, R., Hosoya, S., Kuroyanagi, M., Suzuki, S., Suetake, H., Tasumi, S., Suzuki, Y., Miyadai, T., Kikuchi, K., 2014. An efficient molecular technique for sexing tiger pufferfish (fugu) and the occurrence of sex reversal in a hatchery population. Fish. Sci. 80, 933–942. 10.1007/s12562-014-0768-0

Miao, M., Li, S., Yuan, J., Liu, P., Fang, X., Zhang, C., Zhang, X., Li, F., 2023. CRISPR/Cas9-mediated gene mutation of EcIAG leads to sex reversal in the male ridgetail white prawn *Exopalaemon carinicauda*. Front. Endocrinol. 14. 10.3389/fendo.2023.1266641

Ministry of Agriculture, Forestry and Fisheries of Japan. (2024). Fisheries and Aquaculture Production Statistics, 2022. MAFF. https://www.e-stat.go.jp/stat-search/files?tclass=000001214460&cycle=7&year=20220

Moss, D.R., Moss, S.M., 2006. Effects of gender and size on feed acquisition in the pacific white shrimp *Litopenaeus vannamei*. J. World Aquac. Soc. 37, 161–167. 10.1111/j.1749-7345.2006.00022.x

Nagamine, C., Knight, A.W., Maggenti, A., Paxman, G., 1980. Effects of androgenic gland ablation on male primary and secondary sexual characteristics in the Malaysian prawn, *Macrobrachium rosenbergii* (de Man) (Decapoda, Palaemonidae), with first evidence of induced feminization in a nonhermaphroditic decapod. Gen. Comp. Endocrinol. 41, 423–441. 10.1016/0016-6480(80)90048-9

Nakamura, K., Matsuzaki, N., Yonekura, K.-I., 1992. Organogenesis of genital organs and androgenic gland in the kuruma prawn. Nippon Suisan Gakkai Shi 58, 2261–2267. 10.2331/suisan.58.2261

Omoto, N., Maebayashi, M., Mitsuhashi, E., Yoshitomi, K., Adachi, S., Yamauchi, K., 2002. Effects of estradiol-17β and 17α-methyltestosterone on gonadal sex differentiation in the F2 hybrid sturgeon, the bester. Fish. Sci. 68, 1047–1054. 10.1046/j.1444-2906.2002.00531.x

Pérez Farfante, I., Kensley, B.F., 1997. Penaeoid and sergestoid shrimps and prawns of the world. Keys and diagnoses for the families and genera. Muséum national d’Histoire naturelle, Paris.

Porter, M.L., Pérez-Losada, M., Crandall, K.A., 2005. Model-based multi-locus estimation of decapod phylogeny and divergence times. Mol. Phylogenet. Evol. 37, 355–369. 10.1016/j.ympev.2005.06.021

Posit team, 2023. RStudio: Integrated Development Environment for R. Posit Software, PBC, Boston, MA. Available at: https://posit.co

R Core Team, 2022. R: A language and environment for statistical computing. R Foundation 369 for Statistical Computing, Vienna, Austria. URL https://www.R-project.org/.

Rashid, H., Kitano, H., Hoon Lee, K., Nii, S., Shigematsu, T., Kadomura, K., Yamaguchi, A., Matsuyama, M., 2007. Fugu (*Takifugu rubripes*) sexual differentiation: CYP19 regulation and aromatase inhibitor induced testicular development. Sex Dev. 1, 311–322. 10.1159/000108935

Sagi, A., Cohen, D., 1990. Growth, maturation and progeny of sex-reversed *Macrobrachium rosenbergii* males. World Aquac. 21, 87–90.

Sagi, A., Ra’anan, Z., Cohen, D., Wax, Y., 1986. Production of *Macrobrachium rosenbergii* in monosex populations: Yield characteristics under intensive monoculture conditions in cages. Aquac. 51, 265–275. 10.1016/0044-8486(86)90318-2

Schindelin, J., Arganda-Carreras, I., Frise, E., Kaynig, V., Longair, M., Pietzsch, T., Preibisch, S., Rueden, C., Saalfeld, S., Schmid, B., Tinevez, J.-Y., White, D.J., Hartenstein, V., Eliceiri, K., Tomancak, P., Cardona, A., 2012. Fiji: an open-source platform for biological-image analysis. Nat. Methods. 9, 676–682. 10.1038/nmeth.2019

Schneider, C.A., Rasband, W.S., Eliceiri, K.W., 2012. NIH Image to ImageJ: 25 years of image analysis. Nat. Methods. 9, 671–675. 10.1038/nmeth.2089

Taketomi, Y., Nishikawa, S., 1996. Implantation of androgenic glands into immature female crayfish, *Procambarus Clarkii*, with masculinization of sexual characteristics. J. Crust. Biol. 16, 232–239. 10.1163/193724096X00027

Tropea, C., Hermida, G.N., López Greco, L.S., 2011. Effects of androgenic gland ablation on growth and reproductive parameters of *Cherax quadricarinatus* males (Parastacidae, Decapoda). Gen. Comp. Endocrinol. 174, 211–218. 10.1016/j.ygcen.2011.08.023

Tsutsui, N., Yamane, F., Kakinuma, M., Yoshimatsu, T., 2022. Multiple insulin-like peptides in the gonads of the kuruma prawn *Marsupenaeus japonicus*. Fish. Sci. 88, 387–396. 10.1007/s12562-022-01596-z

Vega-Alpízar, J.L., Alfaro-Montoya, J., Hernández-Noguera, L., Umaña-Castro, R., Aflalo, E.D., Sagi, A., 2017. Implant recognition and gender expression following ampoule-androgenic gland implantation in *Litopenaeus vannamei* females (Penaeidae). Aquac. 468, 471–480. 10.1016/j.aquaculture.2016.11.007

Ventura, T., Manor, R., Aflalo, E.D., Weil, S., Raviv, S., Glazer, L., Sagi, A., 2009. Temporal silencing of an androgenic gland-specific insulin-like gene affecting phenotypical gender differences and spermatogenesis. Endocrinol. 150, 1278–1286. 10.1210/en.2008-0906

Ventura, T., Manor, R., Aflalo, E.D., Weil, S., Rosen, O., Sagi, A., 2012. Timing sexual differentiation: Full functional sex reversal achieved through silencing of a single insulin-like gene in the prawn, *Macrobrachium rosenbergii*. Biol. Reprod. 86, 1–6. 10.1095/biolreprod.111.097261

