## Supplementary material for "Insulin-like androgenic gland factor regulates the development of internal and external secondary sexual characteristics in juvenile male kuruma prawn *Marsupenaeus japonicus*": Figure S1

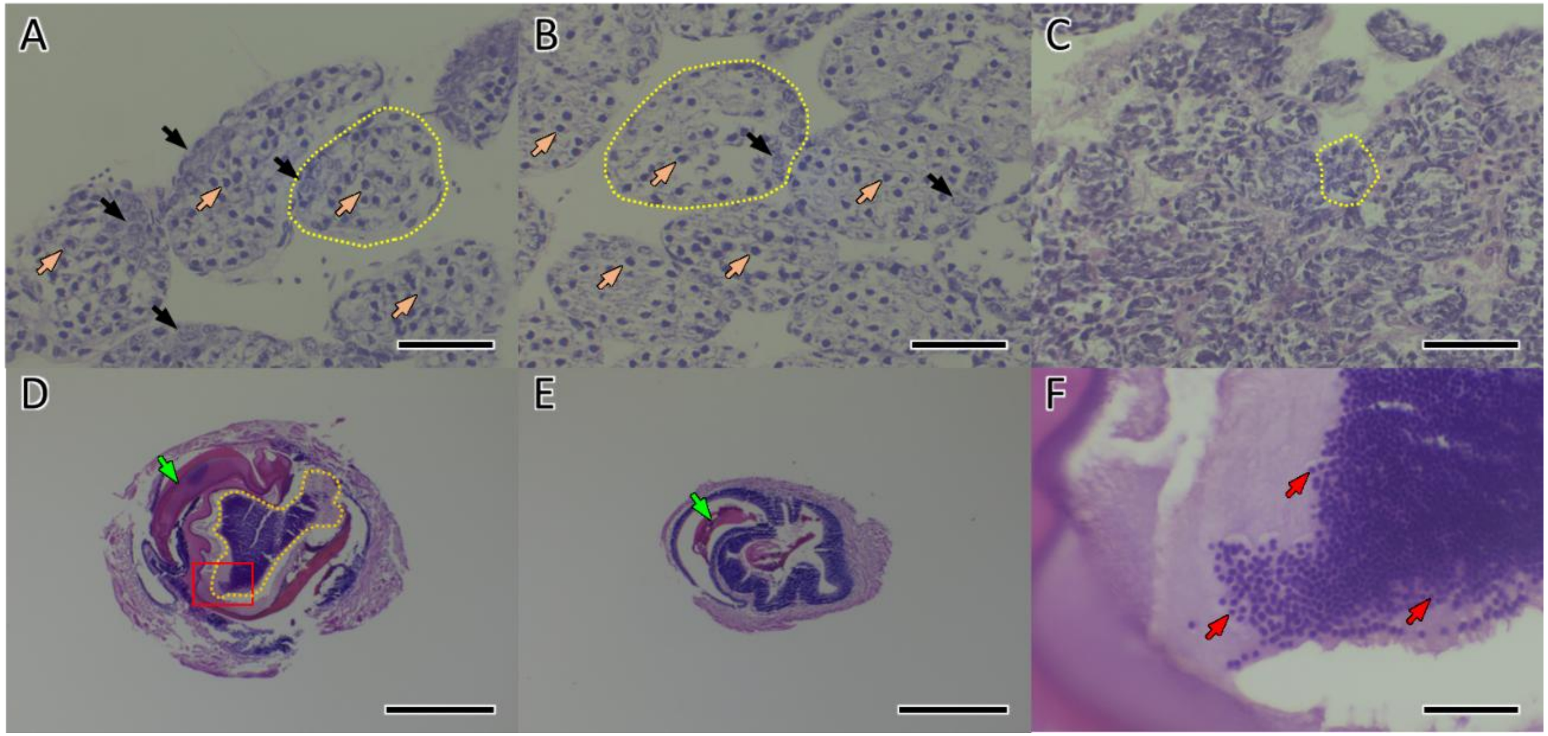


**Figure S1.** Cross-sections of the testis and SV. Testicular sections (A-C) and SV sections proximal to the testis (D-F) are shown. (A), (D), and (F) represent the dsEGFP group at week 9, (B) and (E) represent the dsIAG group at week 9, and (C) represents the intact initial group at week 0. (F) shows a magnified view of the spermatophore region outlined by the red square in (D), as indicated by the red square outlines, where mature spermatozoa are observed. Black arrows indicate spermatogonia, orange arrows indicate primary spermatocytes, green arrows indicate appendages of spermatophores, and red arrows indicate spermatozoa. A typical testicular seminiferous tubule is outlined by yellow dotted lines. The sperm chamber area is outlined with orange dotted lines. Scale bars: 50 µm (A, B, C, F); 500 µm (D, E). Although further examination is needed, appendages of spermatophores were observed in both groups, however, normally developed spermatophores were observed only in the dsEGFP prawns (C, D).
